# Light-dependent inhibition of clathrin-mediated endocytosis in yeast

**DOI:** 10.1101/2021.04.01.432428

**Authors:** Davia Prischich, Javier Encinar del Dedo, Maria Cambra, Judit Prat, Nuria Camarero, Laura Nevola, Andrés Martín-Quirós, Elena Rebollo, Ernest Giralt, María Isabel Geli, Pau Gorostiza

## Abstract

Clathrin-mediated endocytosis (CME) is an essential cellular process, which is evolutionarily conserved among eukaryotes. Yeast constitutes a powerful genetic model to dissect the complex endocytic machinery, yet there is a lack of pharmacological agents that could complement genetics in selectively and reversibly interfere with CME in these organisms. TL2 is a light-regulated peptide inhibitor that targets the AP2/β-arrestin interaction and that can photocontrol CME with high spatiotemporal precision in mammalian cells. Here, we study endocytic protein dynamics by live-cell imaging of the fluorescently tagged coat-associated protein Sla1-GFP and demonstrate that TL2 retains its inhibitory activity in *S. cerevisiae* spheroplasts, thus providing a unique tool for acute and reversible CME modulation in yeast.

## Introduction

Clathrin-mediated endocytosis (CME) is one of the major entry routes for extracellular material and plasma membrane (PM) into eukaryotic cells. The process is essential to internalize nutrients, to maintain PM homeostasis, and to regulate receptor turnover and signalling, among other cellular functions. Endocytic vesicles are also employed by pathogenic toxins,^1–3^ bacteria,^4^ and viruses^5^ (including SARS-CoV-2)^6,7^ to reach intracellular compartments, whereas dysfunctions in the pathway can result in several diseases including cancer, myopathies, and neuropathies^8^. The formation of an endocytic vesicle relies on the sequential and finely coordinated assembly of more than 60 molecular actors which are conserved between yeast and mammalian cells. Thus, *Saccharomyces cerevisiae* provides a convenient model system to study the process, delivering findings that can often be extrapolated to higher eukaryotes^9–11^. Yeast amenability to genetic manipulation, together with advances of live-cell fluorescence microscopy and time resolved electron microscopy, has allowed to examine CME dynamics with great spatiotemporal resolution, and couple the recruitment of particular proteins to specific membrane deformation steps^12–17^.

Several strategies can be used to selectively interfere with CME in order to study the role of specific interactions between endocytic proteins and to decipher the impact of CME on particular cellular processes. Classic genetic manipulation is time consuming and not straightforward, and it can result in compensatory cellular responses that complicate the interpretation of the results. In addition, loss-of-function mutations may result in non-viable strains or display pleiotropic phenotypes when the proteins involved have multiple functions. Reciprocally, genetic redundancy might hinder uncovering of *bona fide* protein functions. These limitations have prompted the search for chemical and pharmacological agents that can specifically, acutely, and reversibly manipulate CME^18^. However, only few chemical tools are known to effectively and specifically inhibit CME, and none of them have been reported to be effective in yeast (**Fig. 1A**)^19,20^. To date, the actin polymerization inhibitor latrunculin A and the sterol sequestering filipin have been effectively used to manipulate endocytic uptake in yeast, but they have pleiotropic effects in many other cellular functions^19,21^. Among the reasons for this shortage of molecular tools is that the endocytic machinery is mainly driven by protein-protein interactions, which involve large contact surfaces that are difficult to tackle with small molecules^22^. Besides, most of these agents display reduced specificity and can affect other endocytic pathways or perturbate other cellular processes^23^.

**Fig. 1.**
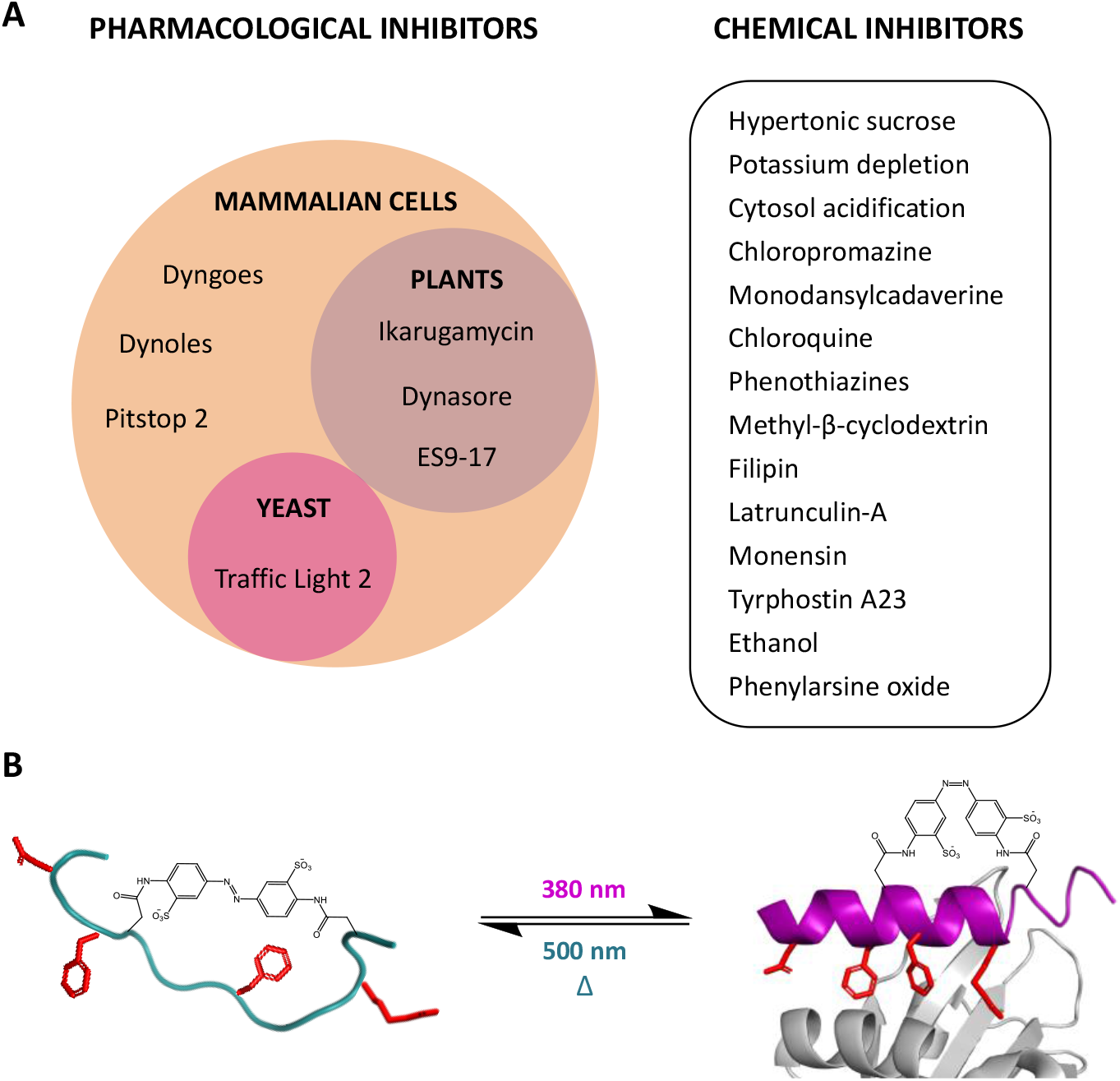
**A)** Overview of clathrin-mediated endocytosis (CME) inhibitors. On the left, pharmacological agents targeting specific endocytic proteins classified according to their biological activity in mammalian cells, plants, and yeast. On the right, chemicals that perturbate CME via a pleiotropic mechanism of action. Limits and advantages of each approach have been covered in the literature^18,20,23–25^. **B)** Traffic Light 2 (TL2) and its structural alterations produced by exposure to light. Upon illumination with UV light (380 nm), the crosslinker photoisomerizes from *trans* (dark-adapted conformation) to *cis*, thus promoting a helical structure (on the right, in purple) and the correct orientation of those residues (in red) responsible for the interaction with AP2 (in grey). Exposure to green light (500 nm) or spontaneous thermal relaxation revert the crosslinker to its stable conformation, thus causing partial unfolding of the peptide and loss of activity (on the left, in green).

Thus, it would be very useful to identify a pharmacological agent capable of interfering with CME in a specific, acute, and reversible way in yeast, to take advantage of this well-defined model for the study of endocytosis. Previously, Nevola *et al*. developed two Traffic Lights (TLs), photoswitchable inhibitors of CME^26^. TLs are based on the structure of the human β-arrestin C-terminal peptide (BAP-long), which binds to the appendage of the β2-adaptin subunit of AP2, the major endocytic clathrin adaptor in mammalian cells^27,28^. Light regulation was introduced by stapling the structures with a photoresponsive bridge that upon *trans-cis* isomerization can modulate the helical content, and therefore, the inhibitory activity of the peptides^29,30^. When tested in mammalian cells, both TLs blocked CME in a light-regulated manner. Between the two, Traffic Light 2 (TL2) proved particularly attractive as it is inert under dark-adapted conditions (*trans* isomer), but it strongly reduces CME trafficking upon activation with (ultra)violet light (*cis* isomer) (**Fig. 1B**). Here, we demonstrate that the TL2 peptide designed for mammalian proteins retains its activity in yeast and provides a powerful tool to dynamically regulate CME. In contrast to small molecules, its peptide nature offers a higher degree of specificity to the mode of action, while the photoswitchable crosslinker enables both the spatiotemporal control of inhibition with light, and the permeability of the compound through the PM.^26^

## Results

Although some peptides have been reported to penetrate the fungal cell-wall, this remains a prohibitive barrier to most macromolecules ^31^. Since TL2 targets an intracellular protein-protein interaction, we first tested whether it could be internalized in yeast using an analogue labelled with carboxy-fluorescein (CF-TL2) on its N-terminus. *S. cerevisiae* and *S. pombe* were incubated in the presence of different peptide concentrations (dark-inactive TL2), but in no case, signs of internal fluorescence were detected, suggesting that cells were unable to take up the peptide or this was quickly extruded (**Fig. S1**). To evaluate if the cell wall might interfere with the peptide uptake, we proceeded to its enzymatic digestion with lyticase before exposing the cells to the fluorescent peptide. To prevent cell lysis, spheroplasts were maintained in media supplemented with 0.7 M sorbitol for osmotic support. A styryl dye, FM4-64 (16 µM), was used to selectively stain vacuole membranes with red fluorescence^32^. This allowed us to confirm that cell integrity, viability, and endocytic activity were maintained after the spheroplasting process. After co-incubation in the presence of fluorescently labelled and dark-adapted inactive CF-TL2, a specific signal corresponding to the peptide was detected by fluorescence inverted optical microscopy in small cortical patches and bright bigger internal structures, as well as in the vacuole, whose limiting membrane was stained with FM4-64. In addition, a weaker signal in the cytosol was detectable. The data thus indicated that under these conditions CF-TL2 was most likely internalized into the cells via an endocytic pathway from where it could be translocated into the cytosol.

To investigate the effect of light-activated TL2 on endocytosis, we followed the cortical dynamics of the GFP-tagged coat-associated protein Sla1-GFP. The yeast clathrin adaptor Sla1 is a component of the clathrin coat, whose spatiotemporal dynamics at endocytic sites has been well-characterized^12,13,33^.

Spheroplasts of *S. cerevisiae* expressing Sla1-GFP were treated with either pre-activated TL2 (380 nm) or with vehicle, and cortical patches were live-imaged after 30 minutes incubation. As shown by kymographs generated from 120 seconds time-lapse movies, light-activated TL2 significantly extended the permanence of the GFP-labelled endocytic patches on the PM (**Fig. 3A**). This effect was dose-, time-, and light-dependent (**Fig. 3B-D**). Inhibition was observed between 50-100 µM and 20-30 min incubation (**Fig. 3B** and **3D**). As light-induced changes are observable close to the IC_50_ and far from saturating doses,^26^ we chose 100 µM and 20 min to quantify light dependency. As reported in mammalian cells,^26^ dark-adapted TL2 did not significantly alter CME, thus confirming that the peptide is also constitutively inert in yeast, and can stall CME upon UV-induced photoisomerization (**Fig. 3C** and **Fig. S2**).

**Fig. 2.**
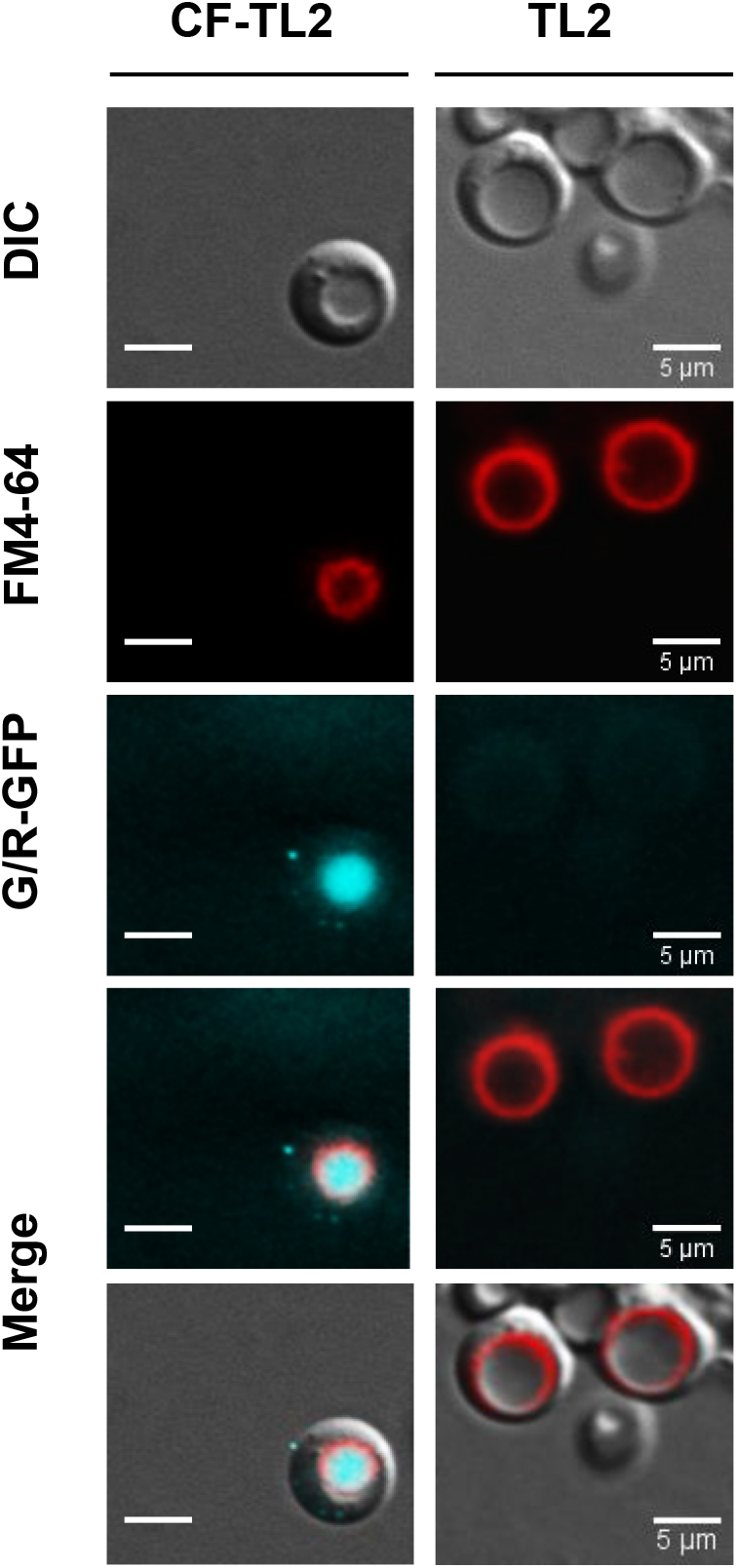
Uptake and intracellular distribution of carboxy-fluorescein conjugated Traffic Light 2 (CF-TL 2) in *S. cerevisiae* spheroplasts as assessed by live-cell fluorescence microscopy. Differential Interference Contrast (DIC) (upper panels) and fluorescence micrographs of *S. cerevisiae* cells loaded for 5 minutes at 25°C with FM4-64 (16 µM), washed and incubated for 40 minutes with 30 µM of either dark-adapted (inactive) carboxy-fluorescein conjugated TL-2 (CF-TL2) or TL2 as control. The individual channels and the merged images of the fluorescein and FM4-64 channels, and the DIC, fluorescein and FM4-64 channels are shown.

**Fig. 3.**
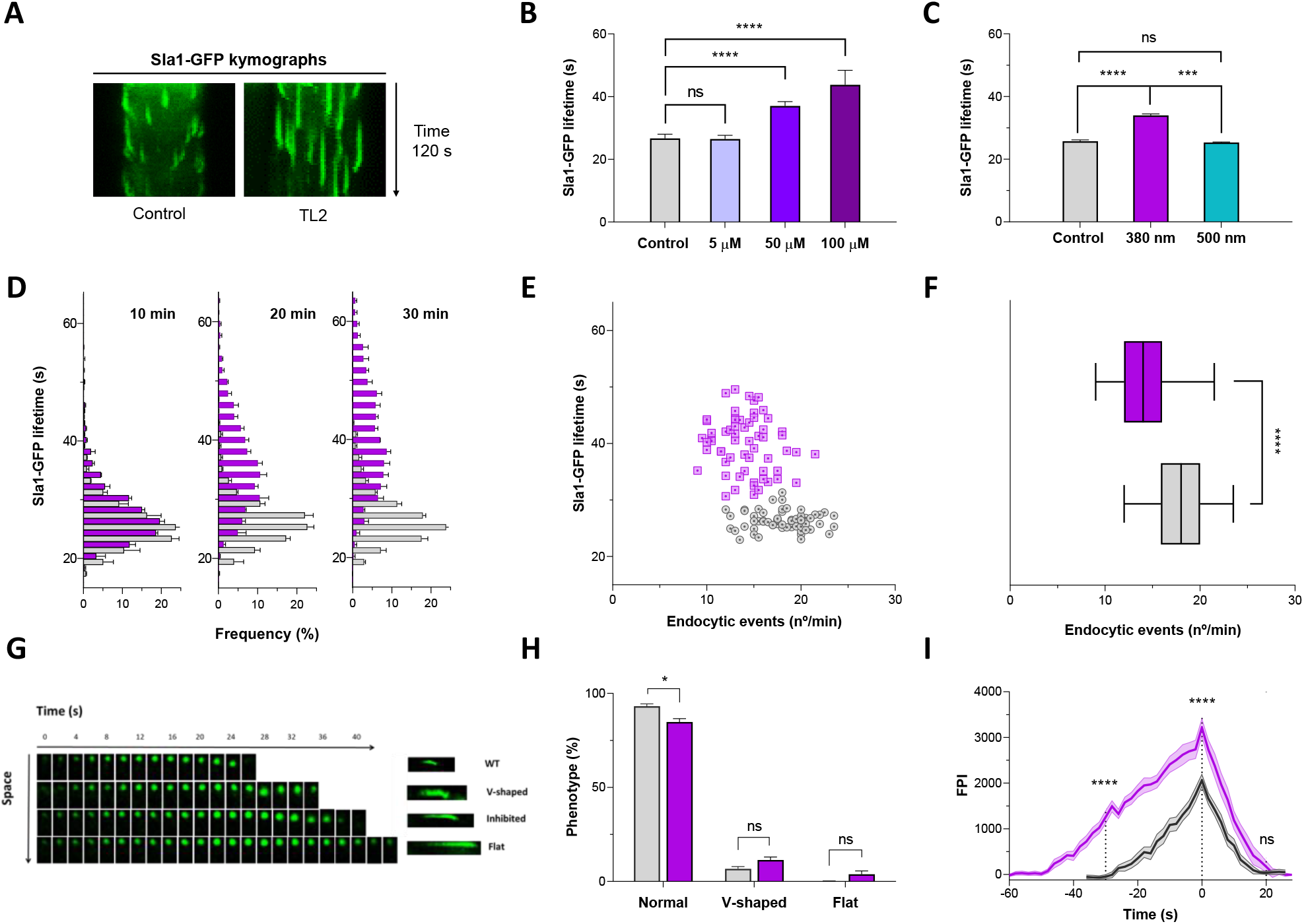
TL2 inhibits CME in yeast in a light-dependent way. **A)** Fluorescence kymographs of Sla1-GFP cortical patches from 120 s time-lapse movies of *S. cerevisiae* spheroplasts incubated with 100 µM light-activated TL2 (illuminated at 380 nm) (right) or mock-treated (left, control). **B)** Mean ± SEM life span of Sla1-GFP cortical patches of yeast spheroplasts either mock-treated (control) or treated with the indicated concentrations of UV-activated TL2 for 30 min. **C)** Mean ± SEM life span of Sla1-GFP cortical patches of yeast either mock-treated (control) or incubated for 20 min with 100 µM of UV-activated TL2 or with the inactive (dark-adapted or illuminated at 500 nm) peptide. **D)** Graphs showing the progressive change in the lifetime distribution of endocytic Sla1-GFP patches incubated either with vehicle or with pre-activated TL2 (100 µM) for 10, 20, and 30 min. **E)** Graph showing the TL2 effects on initiation and maturation of CME sites. Each point represents the average lifespan of cortical Sla1-GFP patches and the number of endocytic *foci* counted per minute in a given cell, 30 min upon incubation with 100 µM UV-activated TL2 (purple) or mock treated (grey). At least 60 cells from 3 different experiments were analysed per conditions. **F)** Graph representing with box and whisker (min to max) the number of endocytic events per cell per minute, in cells incubated for 30 min with 100 µM UV-activated TL2 (purple) or mock-treated (grey). **G)** Time-lapse images of Sla1-GFP cortical patches representing the different dynamics observed under our experimental conditions. Patches were defined as wild-type (WT) when internalization and scission occurred within the described average lifetime for Sla1-GFP. Patches that moved inward and then bounced back towards the PM were classified as v-shaped, independently of their life span. Sla1-GFP dynamics were considered delayed when internalization and scission occurred normally but their cortical life span was extended. Patches were considered flat when no inward movement of the patch was detected. **H)** Frequency plots of patch internalization dynamics, grouped as normal, flat, and v-shaped kymographs (as defined in **G**), in spheroplasts treated with vehicle or the UV-activated peptide (100 µM, 30 min). Classification refers only to the internalization stage not considering the whole duration of the process. Thus, normal patches include both WT and delayed phenotypes. **I)** Comparison of the average fluorescence intensity over time traces for cortical Sla1-GFP patches of spheroplasts treated either with vehicle or with pre-activated TL2 (100 µM, 30 min). Traces were aligned on the fluorescence maximum (time 0). Data are mean ± SEM represented by the shading area. A total of 60 patches from 3 different experiments were analysed per condition. Statistical significance was analysed at time -30, 0, and 20 s. Unless differently stated, statistical differences were determined by Student’s *t* test (n.s., not significant; *, *p*-value<0.05; ***, *p*-value<0.01; ****, *p*-value<0.001). For **B, C, D, F**, and **H**, at least 200 Sla1-GFP patches in at least 20 different cells were analysed per condition. The experiments were repeated at least 3 times and the results plotted together.

Subsequently, we focused on characterizing the active isomer (UV-illuminated TL2, 100 µM) and the extent of its activity in *S. cerevisiae* spheroplasts. Live-cell imaging at different incubation times revealed how TL2 progressively delayed Sla1-GFP internalization resulting in both a shift and a broadening of the patches lifetime distribution when compared to vehicle (**Fig. 3D**). The onset of CME inhibition was consistently observed after 20 minutes, which suggested that the peptide needed to be taken up by endocytosis before translocation to the cytosol. To investigate if clathrin coat assembly, and not only its kinetics, might also be affected by the activated TL2, we quantified both the average Sla1-GFP lifetime, as well as the number of endocytic events per minute after 30 min treatment. Upon plotting these two parameters for each cell, pre-activated TL2- and vehicle-treated samples distributed into two markedly different populations (**Fig. 3E**). CME inhibition associated with a moderate, but significant reduction in the number of endocytic *foci* (**Fig. 3F**), which were 22% less frequent than in the control. We also investigated if the treatment affected later stages of CME (**Fig. 3G-I**). In particular, patches were analysed for their phenotypes and classified according to their internalization patterns (**Fig. 3G**). Normally, the endocytic coat moves into the cytosol as the endocytic invagination grows, and it disassembles as or shortly after the vesicles detach from the PM. In mutants unable to invaginate the plasma membrane though, the endocytic coat remains at the PM (“flat” phenotype). Finally, in mutants affecting the final scission event an initial inward movement of the coat can be observed, but then it bounces back towards the PM (“v-shaped” phenotype) (**Fig. 3G**)^34–37^. TL2-treatment caused marginal changes in the percentage of flat or v-shaped phenotypes compared to vehicle, thus indicating that the peptide does not affect membrane invagination or scission but rather coat maturation and/or cargo loading (**Fig. 3H**). Finally, we analysed the assembly and disassembly dynamics of the Sla1-GFP cortical patches in cells either mock-treated or treated with activated TL2. Sla1-GFP fluorescence intensity plots as a function of time were aligned at their peak intensity (set as time 0) (**Fig. 3I**). Interestingly, while the assembly and disassembly rates appeared similar in both samples, TL2 produced a very significant increase of the maximum fluorescence signal at t=0, revealing that higher levels of Sla1 are required to sustain endocytic budding or that the coats grow larger in the presence of TL2, before initiating membrane invagination (**Fig. 3I**).

## Discussion

We have shown that the photoswitchable peptide TL2 designed from the sequence and structure of human β-arrestin C-terminus can be effectively used to photocontrol CME in yeast. As observed in mammalian cells, TL2 can cross the yeast membranes in minutes and accumulate at concentrations sufficient to inhibit CME. We interpret that pre-illuminated (*cis*, helical) TL2 binds to its target in a conformation that inhibits and slows down clathrin-coated pit (CCP) assembly and internalization, reducing the frequency of events and increasing their duration, thus diminishing the cellular rate of CME. In contrast, the *trans*, less-helical peptide appears inactive in the absence of illumination.

Arrestin-like proteins (ARTs) exist in yeast, which as mammalian arrestins, act as E3 ubiquitin ligase cargo adaptors. Unlike mammalian arrestins though, yeast ARTs lack the specific residues that mediate binding to AP2 and clathrin^38,39^. Interestingly, the yeast epsins Ent1 and Ent2, and the Eps15 homology (EH) domain– containing proteins Ede1 and End3, some of which work as clathrin adaptors for ubiquitinated cargo, all bear sequences similar to the [DE]_n_X_1-2_[FLI]XX[FL]XXXR AP2-binding motif present in mammalian β-arrestins, epsins, and in the TL2 peptide (**Fig. S3**). Therefore, it is likely that the mammalian arrestin function is split in yeast and that TL2 interferes with the clathrin adaptors for ubiquitinated cargo, rather than with the ARTs.

In yeast, AP2 lacks the β2-appendage domain (residues 701-937) that in mammalians interacts with the β-arrestin C-terminus and with TL2 through specific residues located in the protein platform subdomain^26,40^. As the yeast α-appendage shares structural similarities with the mammalian β2-appendage, we investigated if the former could act as binding site of our peptide^27,41^. Adaptin-binding motifs occasionally share similar sequences. In particular, an FXXφ hydrophobic core (where φ indicates a hydrophobic residue) is common to both [DE]_n_X_1-2_[FLI]XX[FL]XXXR and FXXFXXL motifs, which respectively bind the β2- and α-platform subdomains^27,28,41,42^. In both cases, binding is mediated by two hydrophobic sites, which are present in both mammalian appendages and conserved in *S. cerevisiae*. In yeast, a first hydrophobic interaction might be mediated by F907 (mammalian β2-F837) and by I908 and V954, corresponding to mammalian β2-L838 and β2-A877, respectively. The second hydrophobic interaction might be mediated by W911 (β2-W841) and F972 (β2-Y888). In contrast, clusters of polar residues involved in [DE]_n_X_1-2_[FLI]XX[FL]XXXR binding are only found in the mammalian β2-subdomain and in the yeast α-appendage^27,28^. In particular, the basic C-terminus of the peptide could interact with E921 and E982 (β2-E849 and β2-E902 in mammalian), whereas its N-terminal acidic region could interact with R956 (β2-R879 and β2-R834 in mammalians) (**Fig. S4**).

Despite being structurally conserved between lower and higher eukaryotes, AP2 plays a dispensable role in yeast^43,44^. So far, no endocytic phenotype, apart from resistance to the killer toxin K28, has been associated with its knockout, hence it has been suggested that AP2 might act as an adaptor complex for cargoes that are dispensable to yeast viability^45^. Alternatively, the limited phenotypic consequences of deleting AP2 may be due to redundancy of adaptors and/or rerouting to other endocytic pathways. In contrast to genetic manipulation of the AP2 function in yeast, the acute (photo)pharmacological inhibition shown here suggests that AP2 has an important role in CME.

We propose that, once the adaptor complex is recruited into the nascent CCP, TL2 can competitively interfere with protein-protein interactions necessary to advance through the internalization process. Upon binding to the AP2 appendages, TL2 mimics endogenous accessory proteins (clathrin-associated sorting proteins, CLASPs) and hinders progression maybe through the activation of regulatory checkpoints that impede vesicle internalization of incorrectly loaded cargoes^39,46^. Hence, we hypothesize that TL2 acts as a dominant negative peptide exhibiting the AP2-binding motif, but lacking domains essential to regulate endocytic progression. This is supported by the fact that in mammalian models the overexpression of adaptin-binding motifs inhibits CME^42,47^. Moreover, in *S. cerevisiae* the specific binding sequence of TL2 is conserved in several accessory proteins and the corresponding knockout strains display phenotypes that are reminiscent of those obtained upon treatment with TL2. For example, deletion of End3 leads to an increased persistence of Sla1-GFP at the cell cortex^13^, whereas deletion of Ede1 reduces the frequency of site initiations similarly to the presence of the pre-activated TL2, while shortening the lifetime of Sla1-GFP^13,48,49^. Interestingly, an Ede1(22A) mutant that lacks a phosphoregulatory site exhibits reduced patch density (by a 25%) without altering the lifetime of Sla1-GFP^48^. Thus, light-activated TL2 allows to assess the effect of interfering with the [DE]_n_X_1-2_[FLI]XX[FL]XXLR motif on several endocytic proteins and it results in a combination of phenotypes otherwise only appreciable by performing time-consuming, multiple mutations.

## Conclusion

In summary, we have characterized the TL2 activity in *S. cerevisiae* spheroplasts and proved that its ability to control CME with light spans throughout evolution. To the best of our knowledge, no other means to achieve specific pharmacological inhibition of CME have been described to be effective in yeast. While latrunculin A displays a dramatic effect over endocytic vesicle internalization, its inhibitory activity over actin-polymerization interferes with numerous other cellular functions. This, so far, has impeded to understand the contribution of CME to those processes, such as cytokinesis and cell migration, in which the actin role is crucial. In light of these considerations, we believe that TL2 provides a unique tool for the scientific community to gain deeper understandings over the function and regulation of CME. This photoswitchable inhibitor opens the way to intriguing studies where CME could be switch on and off at will using light, in order to synchronize it with other measurements (e.g. microscopy, biochemical reactions) or cellular processes, or obliterate it at specific cellular locations such as the cytokinetic ring or the growth pole.

## Acknowledgements

This research received funding from the European Union Research and Innovation Programme Horizon 2020 - Human Brain Project SG3 (945539), DEEPER (ICT-36-2020-101016787)-, Agency for Management of University and Research Grants/Generalitat de Catalunya (CERCA Programme; 2017-SGR-1442 and 2017-SGR-00465 projects), Fonds Européen de Dévelopement Économique et Régional (FEDER) funds, Ministry of Science and Innovation (Grant PID2019-111493RB-I00), Fundaluce and “la Caixa” foundations (ID 100010434, agreement LCF/PR/HR19/52160010). The project Clúster Emergent del Cervell Humà (CECH, 001-P-001682) is co-financed by the European Union Regional Development Fund within the framework of the ERDF Operational Program of Catalonia 2014-2020 with a grant of 50% of total eligible cost. IBEC and IRB Barcelona are recipient of a Severo Ochoa Award of Excellence from MINECO (Government of Spain). BFU2017-82959-P from MINECO to MIG. D.P. was supported by fellowship BES-2015-072657.

