## Supporting Information for "Light-dependent inhibition of clathrin-mediated endocytosis in yeast"

[1.] Institute for Bioengineering of Catalonia (IBEC), The Barcelona Institute for Science and Technology, Barcelona, Spain
[2.] Centro de Investigación Biomédica en Red – Bioingeniería, Biomateriales y Nanomedicina (CIBER-BBN), Spain
[3.] Department of Cell Biology, Institute for Molecular Biology of Barcelona (IBMB-CSIC), Barcelona, Spain
[4.] Molecular Imaging Platform, Institute for Molecular Biology of Barcelona (IBMB-CSIC), Barcelona, Spain
[5.] Institute for Research in Biomedicine (IRB), The Barcelona Institute for Science and Technology, Barcelona, Spain
[6.] Department of Inorganic and Organic Chemistry, University of Barcelona (UB), Barcelona, Spain
[7.] Catalan Institution for Research and Advanced Studies (ICREA), Barcelona, Spain

‡ Present address: IDP Discovery Pharma, Parc Científic de Barcelona (PCB), Spain
† Present address: Saptools Consulting, Barcelona, Spain

(*) (M.I.G.) (P.G.)

### Materials and Methods

Fluorescently labeled TL2 was synthesized by standard fluoren-9-ylmethoxycarbonyl (Fmoc)-based solid-phase peptide synthesis. Uncrosslinked TL2 was purchased from a custom peptide synthesis service (Ontores Biotechnologies Inc., Hangzhou, CN) and stapled with the BSBCA azobenzene following the procedure described by Nevola *et al.*[1] Lyophilized peptides were reconstituted in 50 mM phosphate-buffered saline (PBS, pH 7.4) at 10x their final concentration. Samples were quantified by measuring the absorption of the azobenzene moiety at 363 nm ($\varepsilon_{363}$ = 24000 $M^{-1}cm^{-1}$). Irradiation was performed using a custom-made multi-LED dual wavelength (380/500 nm) lamp (FC Tecnics, Barcelona, ES). Medium containing TL2 or vehicle was irradiated for at least 10 min before being added to the cells.

In all the assays, protease inhibitors (PIs) were used at the following concentrations: 0.5 mM PMSF, 5 µg/ml Leupeptine, 2,5 µg/ml Antipain, 1 µg/ml Pepstatin and 1 µg/ml Aprotinin. Lyticase from Arthrobacter luteus (≥2,000 units/mg protein; protein ≥20 % by biuret) was purchased from Sigma-Aldrich (Munich, DE) and reconstituted in mQ water. The endocytic marker FM4-64 (Molecular Probes, Eugene, USA) was prepared as a 16 mM stock solution in DMSO. All other chemicals were of analytical grade.

### Strains and Growth Conditions

Yeast strains are listed in **Table S1**. All strains were maintained on petri plates containing the appropriate growth medium supplemented with 2% agar. Rich YPD medium (1% yeast extract, 2% peptone and 2% glucose) was used to culture strains of *Saccharomyces cerevisiae*. *Schizosaccharomyces pombe* cells were grown in YE5S medium (0.5% Difco yeast extract, 3% glucose, 0.025% His, 0.025% Leu and 0.025% Ura). Composition of SDC-Trp medium was as follows: 0.67% yeast nitrogen base, 2% glucose, 0.075% complete supplement mixture (CSM)-Trp. In all the experiments, cells were grown at 25°C under vigorous shaking in Erlenmeyer flasks containing 50 ml of the corresponding growth media. In the morning of the assay, the culture was diluted to an $OD_{600}$ of 0.2 and grown till early log phase (0.45-0.55 $OD_{600}$) before being harvested.

**Table S1.** Yeast strains used in this study

| Strain | Genotype | Source |
|---|---|---|
| SpA64 | *$h^{90}$ leu1-32 ura4-Δ18 his3-d1* | F. Azorín (IBMB-CSIC) |
| SCMIG381 | *Mat***a** *his3 leu2 met15 ura3* | Euroscarf |
| SCMIG933 | *Mat***a** *his3 leu2 ura3 trp1 SLA1-GFP::TRP1* | Fernández-Golbano *et al.* 2014[2] |

### Peptide Uptake in Yeast

From an early log-phase culture, 1.5 ml of cell suspension was harvested and centrifuged for 5 min at 5100 g. The supernatant was replaced with 20 µl of SDC-Trp medium (YE5S medium for *S. pombe*) containing PIs and 100 µM TL2, either fluorescently labeled or not. Cells were incubated at 30°C with gentle shaking and for increasing time intervals (1 hour maximum) before being washed twice and resuspended in 10 µl of medium. Equal volumes of the resulting cell suspension and 1.6% melted agarose (Low Melt Agarose, Bio Rad, Barcelona, ES) were then mixed and an aliquot placed on a microscope slide for imaging.

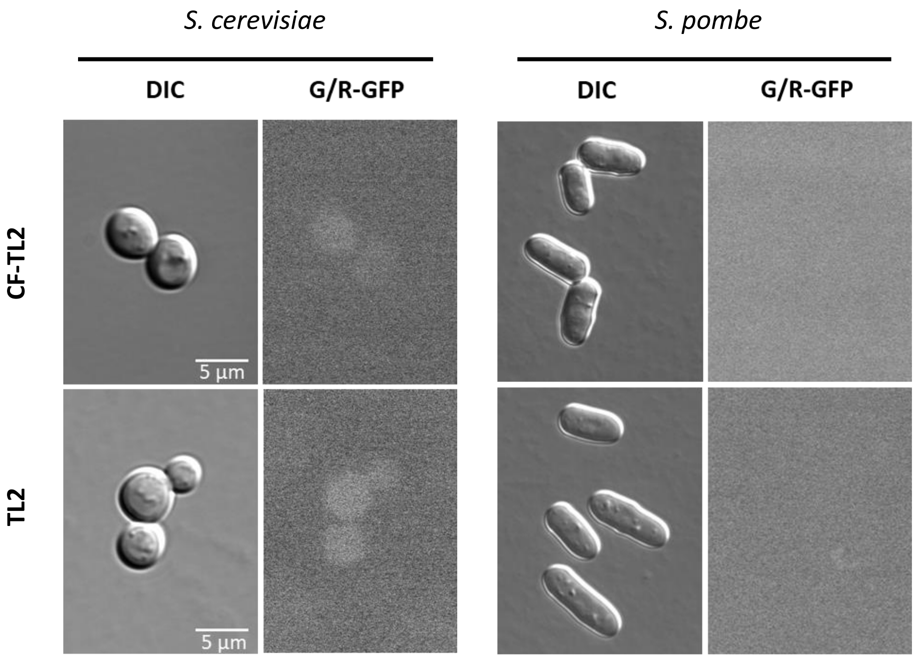

**Figure S1.** Representative differential interference contrast (*DIC*) (left) and fluorescence micrographs of *Saccharomyces cerevisiae* and *Schizosaccharomyces pombe* cells incubated in the presence of 100 µM of either dark-adapted carboxy-fluorescein conjugated Traffic Light 2 (CF-TL2,) or unlabeled TL2 as a control.

### *S. Cerevisiae* Spheroplasts Preparation

Cells were grown in early exponential phase to harvest 15 ml of culture. The suspension was centrifuged for 3 min at 4700 g, washed and the pellet resuspended in half the original volume with 0.1 M Tris HCl (pH 9) supplemented with 10 mM 2-mercaptoethanol. After 10 min incubation at rt and 100 rpm orbital shaking, the culture was washed with 10 ml of spheroplasting buffer (10 mM Tris HCl pH 7.0, 0.7 M sorbitol, 5% glucose, 0.5x YPD-Trp). Cells were collected by centrifugation, resuspended in 1 ml of spheroplasting buffer, and incubated at 30°C and 350 rpm in the presence of 100 u/ml lyticase. After 30 min, the suspension was centrifuged (3 min at 700 g), washed with SDC-Trp supplemented with 0.7 M sorbitol and resuspended in 0.4 ml of the same medium (17-21 $OD_{600}$ U/ml).

### Peptide Uptake in Spheroplasts

Spheroplasts were aliquoted (100 µl per condition, 17-21 $OD_{600}$ U/ml) in Eppendorf tubes and centrifuged for 3 min at 700 g. The supernatant was replaced with 20 µl of uptake buffer (SDC-Trp medium supplemented with 0.7 M sorbitol, 30 µM TL2 – either fluorescently labeled or not – and PIs). After 10 min of pre-incubation at room temperature and 350 rpm, FM4-64 (16 µM final concentration) was added to the suspension and the incubation continued for further 5 min. Spheroplasts were then washed with SDC-Trp medium containing 0.7 M sorbitol, spun down and incubated with 20 µl of fresh uptake buffer for further 40 min. For imaging, cells were washed twice and mounted with low-melting point agarose as previously described.

### Cortical Patch Dynamics

Spheroplasts were aliquoted (100 µl per condition, 17-21 $OD_{600}$ U/ml) in Eppendorf tubes and centrifuged for 3 min at 700 g. The supernatant was replaced with 20 µl of SDC-Trp medium supplemented with 0.7 M sorbitol, 100 µM TL2 (pre-illuminated either at 380 or 500 nm), and PIs. Cells were incubated in the dark at 30°C and 350 rpm, and analysed at different time points (10, 20 and 30 min) as wet mounts in SDC-Trp medium supplemented with 0.7 M sorbitol. To prevent thermal relaxation of pre-activated TL2, vehicle- and peptide-treated samples were exposed to 30 s of UV light after 15 min of incubation.

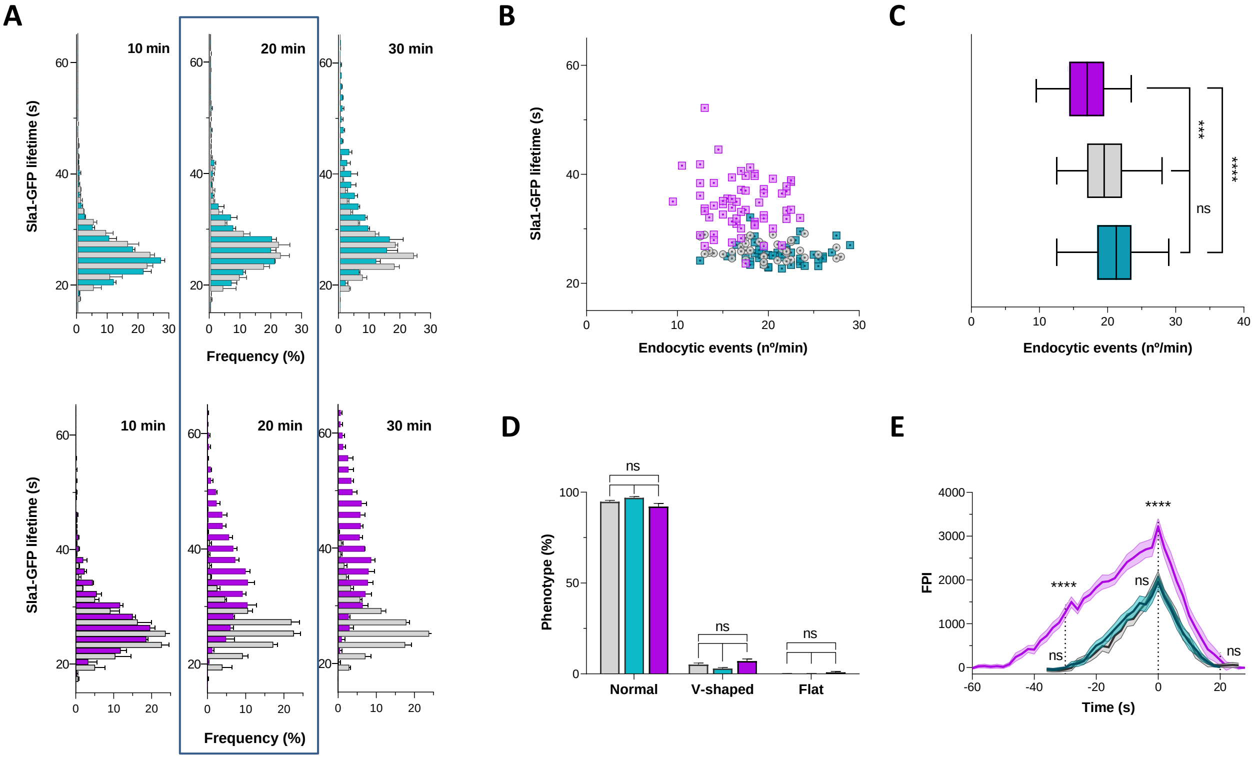

**Figure S2.** Effect of light-activated (in purple) or inactive (in green) TL2 compared to mock treatment (in grey) over CME dynamics. Unless otherwise stated, cells were incubated for 20 min either with vehicle or with 100 µM TL2 (pre-activated or inactive). **A)** Distribution of lifetime frequencies of endocytic Sla1-GFP patches in cells incubated either with vehicle, inactive (dark adapted or illuminated at 500 nm) or active (illuminated at 380 nm) TL2 at 100 µM for 10, 20 and 30 min. The progressive increase in Sla1-GFP lifetime observed with UV-activated TL2 (lower panel) is delayed when spheroplasts are treated with the inactive form of the peptide (upper panel). After 20 min incubation, no difference can be observed between vehicle- or inactive-TL2 treated cells. Data are means ± SEM. **B)** Plot representing the average Sla1-GFP cortical patch lifespan and the number of endocytic events per a given cells per minute in *S. cerevisiae* spheroplasts either mock-treated or incubated with TL2 (pre-activated or inactive). **C)** Plot representing the average number of endocytic events per cell per minute in spheroplasts treated with light-activated or dark-adapted TL2 or treated with vehicle as control. Data is represented with box and whisker (min to max). **D)** Frequency plots comparing patch internalization dynamics in *S. cerevisiae* spheroplasts treated with vehicle, UV-activated or inactive TL2. Kymographs were grouped as normal, flat, and v-shaped on the basis of the phenotype showed by the patch at the internalization stage (see **Fig. 3G**). Data are means ± SEM. **E)** Fluorescence intensity traces as a function of time for endocytic Sla1-GFP cortical patches in spheroplasts treated for 30 min with vehicle or 100 µM TL2 either inactive or pre-activated. Traces were centered on the fluorescence peaks and the point was marked as time 0. Data are fluorescence intensity means ± SEM represented by the shading area (at least 20 patches analyzed per condition and per experiment, $n$=3). Statistical differences were determined at the maximum and the minima of the fluorescence signals (marked by dashed lines). Traces obtained in the presence of inactive TL2 were compared with those from vehicle- and light-activated TL2 treated samples. Unless otherwise stated, statistical differences were determined by Student's $t$ test (n.s., not significant; ***, $p$-value<0.01; ****, $p$-value<0.001). For **A**, **B**, **C**, and **D** at least 200 patches in at least 20 cells were analyzed per condition and per experiment. Experimental $n$ = 2 (inactive TL2) and $n$ = 3 (active TL2 and vehicle).

```
TL2                          DDDIVFECFARQRLCGMKDD
β-arrestin2(380-398)       YATDDDIVFEDFARLRLKG
Epsin1(397-414)            FDTEPD-EFSDFDRLRTAL
Epsin2(451-468)            NGTIKD-DFSEFDNLRTSK

Ent1(364-381)              TGTGID-TFGNVGEARIPA
    (92-109)               VLWCRE-NLYIIKTLKEFR
Ent2(448-465)              TGTGID-TFGNTGEARIPA
    (92-109)               VLWCRE-NFYVIKTLREFR
Ede1(1019-1037)            QSVRDDVELPETLEERDTI
End3(183-201)              PSIDRDPTFYFIHCLRQRN
```

**Figure S3.** The AP2 interaction motif $[DE]_nX_{1-2}$[FLI]XX[FL]XXXR in TL2, human β-arrestin2 and epsins, and possible conserved motifs from the yeast epsin homologues, Ent1 and Ent2, and the Eps15 homology (EH) domain–containing proteins, Ede1 and End3. Hotspots mediating binding to AP2 are highlighted in yellow. Residues in red are conservative substitutions between the mammalian and yeast proteins. Further conserved residues are highlighted in grey.

```
AP2B1 human   GK-MERQVFLATWK---DI-PNENELQFQIKEC-HLNA--DT--VS-
AP2A  yeast   MHLNLAQ-FISRWKTLSDALG-K-EGEYQ-KSGIKLNK--DFRKVET
AP2A1 human    --MAAQDFFQRWKQLS--LPQ-Q--EAQ-KIF-KANHPMDAE-VTK

AP2B1 human   -SKLQNNNVY---T-IAKR----NVEGQDM----LYQSLKLTNGIWI
AP2A  yeast   IS-LEDGLL--LLTQTVKRLGFDIVD-QTSVRSTLFVS-----GI-I
AP2A1 human   A-KLL-GFGSALL---------DNVDPNP----ENFV----GAGI-I

AP2B1 human   LA--E------LRIQPGNP-NYT--LSLKC
AP2A  yeast   HTKSEGNFGCLMKIQY--QVNGTVNVTCKT
AP2A1 human   QTKALQV-GCLLRLEPNAQAQMYR-LTLRT
```

**Figure S4.** Sequence alignment between the human β2-appendage, and the human and yeast α-appendage platform subdomains from AP2. Residues binding to the TL2 interaction motif $[DE]_nX_{1-2}$[FLI]XX[FL]XXXR are highlighted in yellow, conserved substitutions in red. Residues marked in cyan mediate binding to cargo adaptors in the human α-appendage. Additional conserved residues between the three domains are pictured in grey.

### Fluorescence Microscopy

Fluorescence microscopy was performed as described by Del Dedo *et al.*[3] Samples were viewed on a Leica DMI6000 wide-field microscope equipped with a 63x APO (NA=1.4) oil immersion objective. Images were acquired at rt using a Hamamatsu Orca-R2 camera controlled by the Leica Application Suite X (LAS_X) software. Fluorescence was visualized with appropriate filter sets mounted on an ultrafast external wheel and illuminated by a SOLA-SMII white LED light. GFP or carboxyfluorescein and FM4-64 were excited through BP470/40 and BP572/35 filters, respectively. Emitted light was detected with BP521/40 (green) and BP632/62 (red) filters. Movies were acquired at 2 s intervals during 2 min.

### Quantification and Statistical Analysis

ImageJ (NIH, Bethesda, USA)[4] was used to adjust image size, brightness and contrast in figure 2, and to project and quantify Sla1-GFP kymographs for figures 3 and S1. In figure 3I and S1E, fluorescence intensity was plotted after subtracting the relative background intensity with Microsoft Excel. All graphs were created with GraphPad Prism v8.3.1 (GraphPad Software, San Diego, USA). Statistical analysis was performed either with GraphPad Prism or Microsoft Excel.

[1] L. Nevola, A. Martín-Quirõs, K. Eckelt, N. Camarero, S. Tosi, A. Llobet, E. Giralt, P. Gorostiza, *Angew. Chemie - Int. Ed.* **2013**, *52*, 7704–7708.

[2] I. M. Fernández-Golbano, F. Z. Idrissi, J. P. Giblin, B. L. Grosshans, V. Robles, H. Grötsch, M. M. Borrás, M. I. Geli, *Dev. Cell* **2014**, *30*, 746–758.

[3] J. Encinar del Dedo, F. Z. Idrissi, I. M. Fernandez-Golbano, P. Garcia, E. Rebollo, M. K. Krzyzanowski, H. Grötsch, M. I. Geli, *Dev. Cell* **2017**, *43*, 588-602.e6.

[4] C. A. Schneider, W. S. Rasband, K. W. Eliceiri, *Nat. Methods* **2012**, *9*, 671–675.
